# Multistage feedback driven compartmental dynamics of hematopoiesis

**DOI:** 10.1101/2020.09.10.291393

**Authors:** Nathaniel V. Mon Père, Tom Lenaerts, Jorge M. Pacheco, David Dingli

**Affiliations:** Interuniversity Institute of Bioinformatics in Brussels, Université libre de Bruxelles, Brussels, Belgium; Applied Physics group, Department of Physics, Vrije Universiteit Brussel, Brussels, Belgium; Machine Learning Group, Département d’Informatique, Université libre de Bruxelles, Brussels, Belgium; AI lab, Computer Science Department, Vrije Universiteit Brussel, Brussels, Belgium; Centro de Biologia Molecular e Ambiental, Universidade do Minho, 4710–057 Braga, Portugal; Departamento de Matemática e Aplicações, Universidade do Minho, 4710–057 Braga, Portugal; ATP-group, P-2744-016 Porto Salvo, Portugal; Division of Hematology and Department of Molecular Medicine, Mayo Clinic, Rochester, MN, United States of America

## Abstract

Human hematopoiesis is surprisingly resilient to disruptions, providing suitable responses to severe bleeding, long lasting immune activation, and even bone marrow transplants. Still, many blood disorders exist which push the system past its natural plasticity, resulting in abnormalities in the circulating blood. While proper treatment of such diseases can benefit from understanding the underlying cell dynamics, these are non-trivial to predict due to the hematopoietic system’s hierarchical nature and complex feedback networks. To characterize the dynamics following different types of perturbations we investigate a model representing hematopoiesis as a sequence of compartments covering all maturation stages – from stem to mature cells – where feedback regulates cell production to ongoing necessities. We find that a stable response to perturbations requires the simultaneous adaptation of cell differentiation and self-renewal rates, and show that under conditions of continuous disruption – as found in chronic hemolytic states – compartment cell numbers evolve to novel stable states.

## Introduction

Hematopoiesis is the physiological process responsible for the production of all circulating blood cells. This includes the oxygen-carrying erythrocytes and several types of white blood cells associated with the innate and adaptive immune response and platelets. The general mechanism follows a hierarchical architecture, with rare slowly replicating multipotent hematopoietic stem cells (HSCs) seeding more differentiated progenitors that increase in frequency through successive levels of maturation [1–3]. Considering the ubiquitous nature of hematopoietic cell types in the body, it is no surprise that we observe many disorders – often hereditary in origin – related to improper development or problematic behavior in the bone marrow [4].

Detailed experimental studies of various aspects of this process can be found dating back to the previous decennium – spanning topics such as hematopoietic stem cells [5], lineage development [3,6] and signaling pathways [7] – resulting in a qualitatively detailed picture of this architecture. However, from a quantitative viewpoint our collective knowledge is still lacking, in no small part due to the fact that in vivo studies of the bone marrow cell dynamics present numerous challenges. Still, in the past decades a handful of mathematical models of hematopoiesis have been developed [8–19] leading to important insights on several of its properties.

In such a complex multicompartment structure that is influenced by the presence of cytokines, chemokines, hormones, and the local microenvironment, regulatory feedback loops linking such compartments are expected to be present. Such feedback loops likely have important consequences on the overall cell dynamics, especially after the occurrence of perturbations which disrupt normal homeostasis. Understanding the types of behavior which may occur can be important for understanding the progression of hematopoietic diseases which affect blood cell production, concentration, and development. Here we investigate some properties of this class of systems, with a focus on how perturbations to cell numbers can influence the self-renewal and differentiation behavior of the maturing cells. To this end we develop a theoretical model which, starting from the description of hematopoiesis developed by Dingli et al. [11], introduces regulatory feedback mechanisms that allow the system to react to perturbations, for example a loss of cells due to bleeding or hemolysis. In the following, we introduce a general formalism for modeling feedback in linked hierarchical compartments by specifying the requirements of such a coupling without specific knowledge of the coupling function itself. Subsequently, we apply this feedback structure to examine the dynamics of a multistage feedback-driven compartmental model of hematopoiesis, validated using data from a study on erythrocyte dynamics [20]. We study possible dynamic behaviors after a perturbation, and identify under which conditions these occur. Finally we assess how the behavior changes under chronic perturbations.

## Model and Methods

The model of Dingli et al. [11] constitutes our starting point. It describes the maturation process of hematopoietic cells through a fixed number *M* of discrete compartments associated with progressive “levels” of differentiation that all cells traverse before leaving the bone marrow. Within each compartment *j* a cell divides at a predefined rate *η*, where each division is considered symmetric (for simplicity), that is, it gives rise to two identical daughter cells. These are either exact replicas of the parent – with probability 1 – *ϵ_j_* – and thus remain in the current compartment *j,* or have differentiated – with probability *ϵ_j_* – and thus move to the subsequent compartment *j* + 1. Under homeostatic conditions the number of cells in each compartment should remain approximately constant in time, while compartment sizes increase toward maturity at a fixed ratio *N*_*j*+1_/*N_j_* = *η* to accommodate the expansion of a small number of stem cells (*N*_0_, of the order of several hundred for humans) to the daily output of the bone marrow (*N_M_* ≈ 10^11^). This exponential increase is mirrored by the division rates: *r*_*j*+1_/*r_j_* = *ρ*, while the differentiation probability is the same for all compartments: *ϵ_j_* = *ϵ*. Values for these parameters can be derived by fixing the initial and final compartment sizes and division rates, and using the equilibrium requirement *∂_t_N_j_* = 0. (see Supplementary material).

In order to address the coupling between compartments through feedback, we now alter the existing formalism. First, we formally describe both types of division – self-renewal (*j* → *j*) and differentiation (*j* → *j* + 1) – as independent Poisson processes occurring with rates *v_j_* and *s_j_* respectively. It can be shown that this description is equivalent to the original one through the relations *r_j_* = *s_j_* + *v_j_* and *ϵ_j_* = *s_j_*(*s_j_* + *v_j_*)^−1^ (see Supplementary material 2 for a detailed derivation). The cell dynamics are given by the following equations for cell numbers in each compartment *j*:

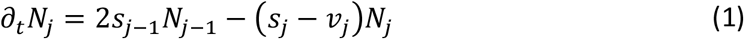

Under homeostatic conditions the system is stable with 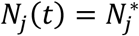, and 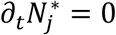, and the division rates are given by their homeostatic values 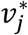 and 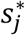. We introduce feedback through sequential coupling between successive compartments, by allowing the rate parameters of each compartment to vary depending on the number of cells in a neighboring downstream compartment. Given a perturbation 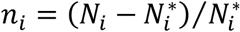 on the cell number in compartment *i,* we are thus looking for non-negative functions *v_j_*(*n_i_*) and *s_j_*(*n_i_*) that produce a negative feedback response – i.e. opposing the sign of *n_i_*,. Furthermore, we assume there is an upper limit to how many divisions a cell can undergo, thus determining upper bounds on *v_j_*(*n_i_*) and *s_j_*(*n_i_*). Naturally, the fact that homeostasis is maintained in the absence of any perturbation implies that 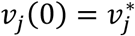 and 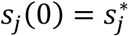.

From the outset, the functions *v_j_*(*n_i_*) and *s_j_*(*n_i_*) are expected to be the solution of a highly non-linear ecological network of various cell types, nutrients, and signaling factors [21]. Here, instead, we look for the simplest functional form that fulfills the requirements above; this leads us to the linear form

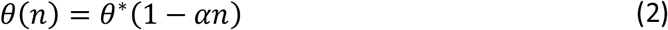

(with the placeholder *θ* = {*v, s*} and *α* > 0 to ensure negative feedback), such that 0 ≤ *θ*(*n*) ≤ *θ_max_* ≡ *k_θ_θ*\* (Fig 1). A smoother version is easy to define by drawing inspiration from classical ecological systems, which mirror the competition for promoting or inhibiting factors among different cell groups [21,22]. In this vein, a logistic function (Fig 1) of the form

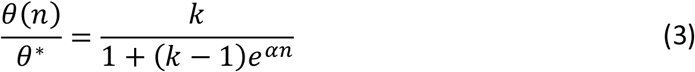

where the parameters *k* and *α* play analogous roles in determining respectively the maximum and the slope, constitutes a natural choice.

**Figure 1.**
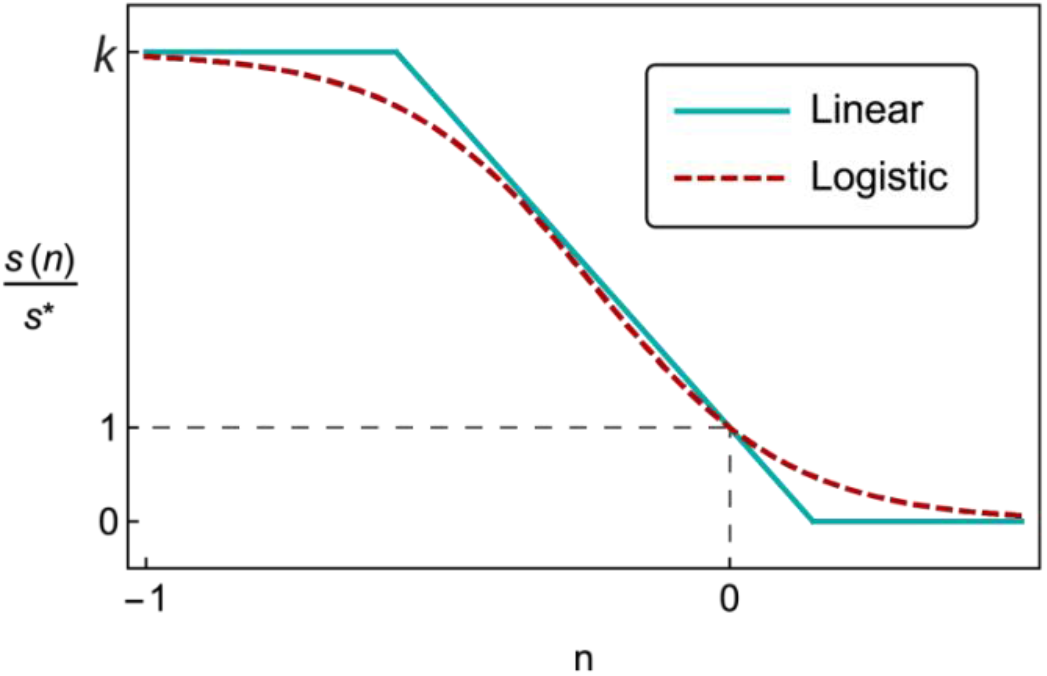
Illustration of linear and logistic differentiation rate functions. *Both are bounded between* 0 *and ks*, and have s*(0) = *s*\*.

While the rate functions defined above provide a useful method for coupling any pair of compartments, modeling the full hematopoietic system requires an interaction network that defines the pairwise connections between compartments. Many complex circuits are possible, and the number of potential interaction combinations (through pairs or higher orders) increases dramatically with the number of compartments.

Here we explore a simple case, in which all compartments are coupled sequentially to their downstream neighbors, so that the rate functions have the form *s_j_*(*n*_*j*+1_) and *v_j_*(*n*_*j*+1_) for all *j*. Given this interaction network, as well as the rate functions and their parameters *α_s_*, *k_s_*, *α_v_*, *k_v_* ∈ ℝ^+^, the solution to (1) for *M* compartments can be obtained numerically through any finite difference method.

## Results

### Sequential coupling elicits three types of behavior

We start by examining the case in which hematopoiesis proceeds under homeostasis when a perturbation occurs in a single compartment. The response in the absence of feedback mechanisms has been studied previously in [23] and can be recovered in this model by fixing the division rates to their homeostatic values: 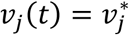 and 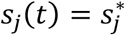. Equation (1) shows that without feedback the compartmental coupling is entirely one-directional and upstream: the dynamics of *N_j_* depends on *N*_*j*−1_ but not on *N*_*j*+1_, meaning that compartments will not respond to disturbances taking place in downstream compartments. Still, when a transient perturbation from equilibrium occurs in a given compartment *j*, the homeostatic equilibrium is eventually restored (Fig 2a), though in the absence of downstream coupling the relaxation time is too long to match real recovery times (see discussion). While all upstream neighbors *j* – *k* remain in homeostatic conditions (*n*_*j*−*k*_ = 0) all downstream *j* + *k* are affected as the perturbation moves successively through these compartments.

**Figure 2.**
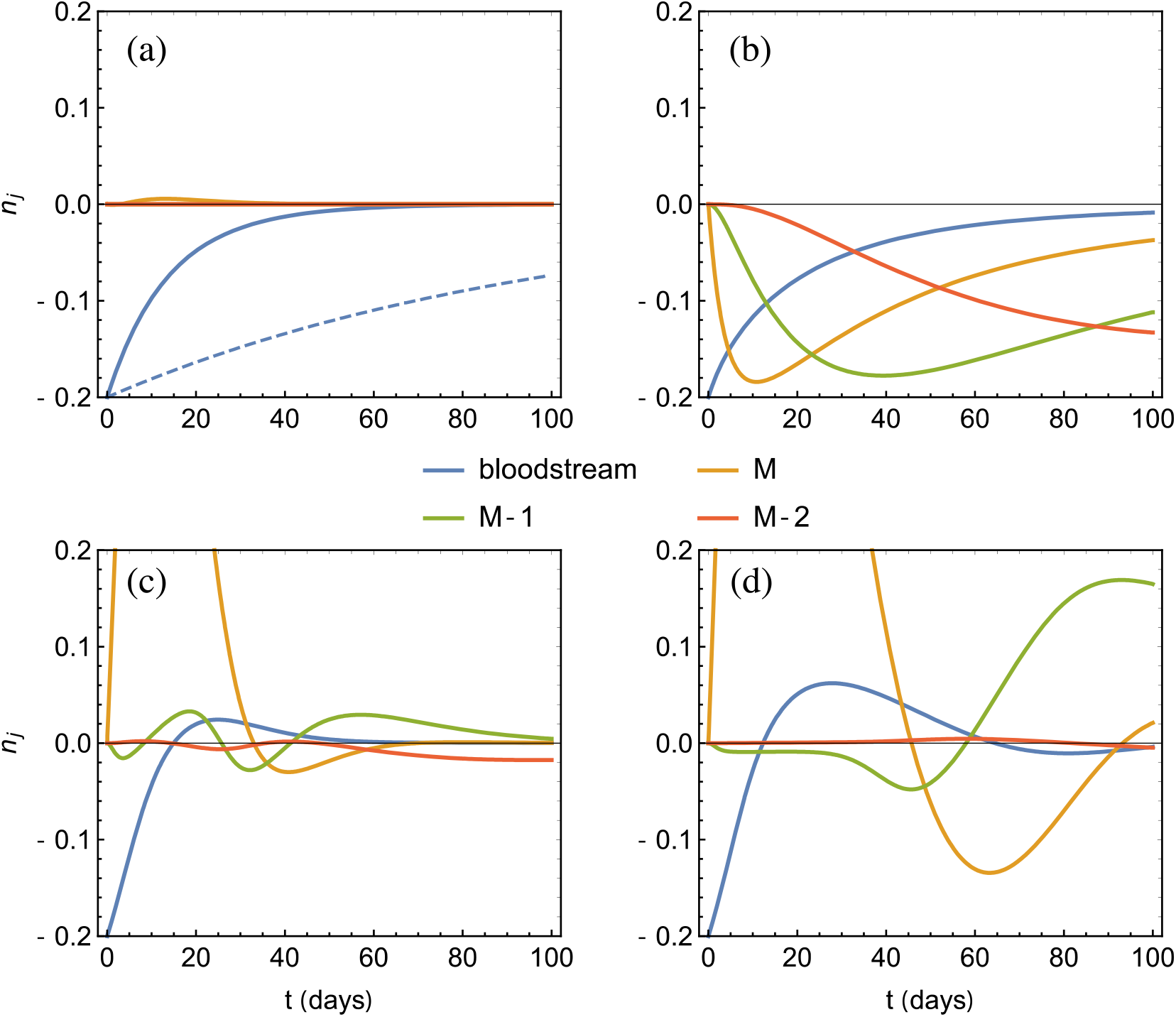
Compartment number dynamics of logistically coupled feedback systems following a sudden loss of cells in the bloodstream. The cell number is expressed in relative perturbation 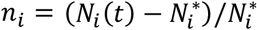, and the bloodstream and final three compartments (M-2, M-1, M) are shown. Parameters k_s_, k_v_, and α_s_ are obtained from parameterization to Hillman et al. [20] (see main text and Fig 3 for details). **(a)** An M=5 compartment model without feedback (dotted line) and with balanced response (∂_n_v ≈ ∂_n_s) feedback (full lines). The response is not entirely without upstream propagation due to the logistic character of the rate functions, for which no perfectly balanced solution (equation (4a)) exists. **(b)** Differentiation driven response (∂_n_s < ∂_n_v) with resulting positive feedback (M=5). **(c)** Self-renewal driven response (∂_n_v < ∂_n_s) with oscillatory behavior (M=5). **(d)** Same as (c) except that M=3.

This behavior will change when feedback – as described above – is introduced: A dependence of *N_j_* on *N*_*j*+1_ is now included and a similar wavelike propagation upstream is now expected. The key components of our model that determine the dynamics following a perturbation are the ratio of the coupling strengths of the differentiation/self-renewal rates *s_j_*(*n*_*j*+1_)/ *v_j_*(*n*_*j*+1_), and the total number of feedback stages in the system. The latter will be discussed in the next section. To understand the former – the effect of the relative coupling strengths – we define the simplest possible network with just a single coupled “pair”, and turn our attention to the state of the system at time *t*_0_> immediately after a perturbation *n*_*j*+1_ is introduced, so that *n_j_* is still 0. Then the dynamics (1) of the reacting compartment *j* can be rewritten as

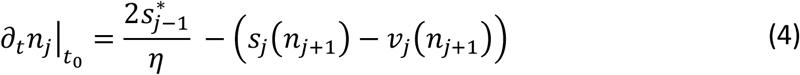

If the outgoing flux is unchanged from the homeostatic case, i.e.

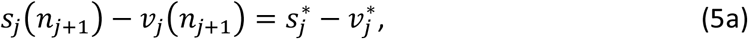

the equilibrium condition is achieved and we have *∂_t_n_j_* = 0. If this remains true for any value of *n*_*j*+1_ then under sequential coupling this condition prevents the perturbation from moving upstream with respect to the first responding compartment, and thus ‘protects’ all upstream compartments from deviating from homeostasis while significantly reducing the time required to return to homeostasis (Fig 2a). For the linear rate functions (2), equation (1) leads to the following condition to ensure that equation (4a) is fulfilled:

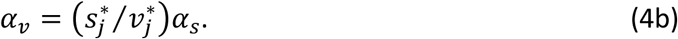

While in general no such solution exists for the logistic functions, we use this relation as a first order approximation and denote 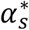 and 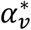 to indicate parameter values which fulfill this requirement.

Whenever equation (4a) is not fulfilled, then *∂_t_n_j_* ≠ 0 and one can expand the rate functions about *n*_*j*+1_ = 0 which, after cancellation of the zeroth order terms (due to the homeostatic condition) gives:

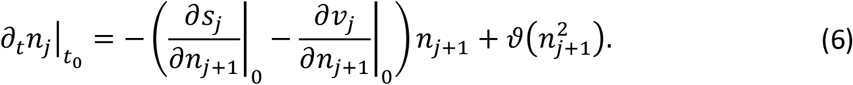

Ignoring higher order terms and recalling that we have required *∂*_*s_j_*_/*∂*_*n*_*j*+1__ < 0 and *∂*_v_j__/*∂*_*n*_*j*+1__ – 0 to ensure negative feedback, we see that the sign of *∂*_*t*_*n_j_*|*t*_0_ can either oppose or match the sign of *n*_*j*+1_, depending on the difference in the brackets: *∂*_*n*_*v* < *∂*_*n*_*s* or *∂*_*n*_*s* < *∂*_*n*_*v* respectively. In the latter case the matching sign means the feedback can actually amplify rather than dampen the perturbation, provided the difference is large enough, since a loss (or excess) of cells would induce further losses (or excesses) in upstream compartments that are required to provide the incoming flux of cells; this in fact corresponds to a positive feedback, as shown in Fig 2b. Conversely, if *∂*_*n*_*v* < *∂*_*n*_*s* we obtain the desired negative feedback regime. Nonetheless, damped oscillations may emerge (Fig 2c) which, if severe, can prolong the time necessary for the system to return to homeostasis. Recalling that s and *v* are strictly decreasing functions, the condition *∂*_*n*_*v* < *∂*_*n*_*s* implies that the rate of self-renewal changes faster with *n* than the rate of differentiation, while *∂*_*n*_*s* < *∂*_*n*_*v* implies the contrary. Thus intuitively we can interpret these conditions as determining whether the response is driven more by increased self-renewal (*∂*_*n*_*v* < *∂*_*n*_*s*), increased differentiation (*∂*_*n*_*s* < *∂*_*n*_*v*), or if the increase is balanced across both processes (*∂*_*n*_*s* = *∂*_*n*_*v*).

### Increasing cell amplification between compartments reduces stability

As previously stated, the number of interacting compartments *M* also influences the overall dynamics. Note that *M* need not necessarily be the same as the number of differentiation stages found through traditional methods such as surface marker identification or transcriptional profiling, as our treatment is flexible enough to loosely describe stages of development which interact through feedback, and thus these compartments may encompass multiple maturation stages found in other models. To ensure a meaningful comparison, we change *M* assuming the same number of cells at the root of the hematopoietic tree and under circulation. Thus, smaller *M* implies larger cell amplification rates between consecutive compartments.

Varying *M* is found to influence the stability of the hematopoietic system with respect to the rate parameters. Indeed, when deviations from the conditions in equations (4a) and (4b) take place, one obtains an increase in amplitude of oscillations with decreasing *M* (Figs 2c and 2d). The origin for this can be seen even when employing the linear coupling function in equation (4) (which is equivalent to keeping only the linear terms in the logistic function): there, the inequality in the first derivative becomes *α_v_*/*α_s_* ≠ *s*\*/*v*\*, meaning that the amplitude of the oscillations is determined by how much *α_v_/α_s_* deviates from *s*/v\**, the ratio of homeostatic division rates. In particular, perturbations on 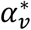 or 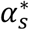 will have a larger impact the smaller this ratio is. Furthermore, it can be shown that *s*/v\** decreases monotonically with increasing cell number amplification *η* between compartments, which in our model is akin to decreasing *M.* In this sense hematopoietic models with lower *M* are less stable under perturbations on the parameters *α_s_* and *α_v_*. It is worth noting here that stability under variation of these parameters forms an important requirement for the system itself and will be discussed in detail later.

### Recovery time as a measure of efficiency

The time for a compartment to recover from a perturbation is an important measure of the efficiency of hematopoiesis, as an expedited recovery can be considered more advantageous for the host. This recovery time is directly determined by the strength of the response to a loss of cells, which the model itself sets little restriction on: The *k* and *α* parameters – respectively determining the maximal increase in divisions and the severity of the perturbation at which this maximal value is reached – can technically (i.e. as long as equation (4a) is fulfilled) be taken arbitrarily high without inducing oscillations or positive feedback. However, in real hematopoiesis one would expect physical limitations to apply to these, such as for example the time and/or resources required for cells to undergo additional divisions.

In addressing the recovery time, we should take all possible recovery types into consideration. Indeed, we should keep in mind that hematopoietic cell numbers fluctuate in time even under homeostatic conditions [4]. Consequently, it is reasonable to assign some range around the model’s equilibrium value within which a compartment can be considered “recovered”. For example, while Fig 2a shows a greatly improved response compared to the feedback-free model, the oscillatory behavior in Fig 2c presents a qualitatively superior result with respect to the recovery time – effectively halved in this scenario – if we consider a compartment to be recovered once it has returned to within approximately 2% of its homeostatic value. Thus a slight emphasis on self-renewal rather than differentiation in the response can be beneficial if the resulting oscillations are small in amplitude. Conversely, while the regime depicted in 2b (emphasis on differentiation) also improves upon the feedback-free model, it is less efficient than that of 2a, as the resulting positive feedback always reduces efficiency (1b).

### Inclusion of feedback allows prediction of erythrocyte dynamics

To evaluate the predictive power of the model we use data from Hillman et al. [20], who study the human bone marrow response to a severe loss of erythrocytes. The authors mark the increase in erythrocyte production as a function of the normal output for different levels of depletion of the hematocrit, noting that the efficiency of the response depends strongly on the amount of iron available to the patient. We can translate the hematocrit measurements to perturbations in our model by taking the ratio of the depleted to the normal value; for example if the patient’s normal hematocrit is 50%, a reduction to 40% would equate to a 20% loss, which is a perturbation in the bloodstream compartment (*B)* of *n_B_* = −0.2. A summary of their findings is shown in Fig 3. We estimate our parameter values by assuming 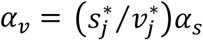 and making *k_s_* = *k_v_* ≡ *k*. For this coupling the dynamics of the perturbed bloodstream compartment can be written as *∂_t_n_B_* = 2*s_M_* – *μ_B_*(1 + *n_B_*), with *μ_B_* the constant loss rate of circulating cells; which is independent of the replication rate function *v_M_* of the preceding compartment. Thus *α_v_* is fixed by the response requirement and only *k* and *α_s_* are free. A least-squares fit of the logistic coupling (3) results in parameter pairs for the three patient cohorts defined by the authors (based on the patients’ body iron stores). The values for the normal patient cohort (*k* = 3.5, *α_s_* = 7.5) are used in Fig 2. Different parameter pairs are found for the other cohorts, with a clear effect being an increase in maximal production factor *κ* for increasing iron availability. This implies that the response relies not only on the severity of the perturbation but on the availability of essential resources as well, so that the parameters *α_s_*, *α_v_*, *κ_s_* and *κ_v_* should in fact depend on other parameters reflecting a dynamic environment. The values for *k* = *k_v_* = *k_s_* found here to range between 3.5 (normal cohort) and 7.6 (hemochromatosis) fit with current knowledge of production rates of mature red blood blood cells, where the highest reported rate increases are 8- to 10-fold the normal rate [20]. For this range of *k* we thus estimate the slope parameter *α_s_* to be in the range of 7.2-11.3, while *α_v_* is then determined by the compartment number through *α_v_* = *α_s_ s*\*/*v*\*.

**Figure 3.**
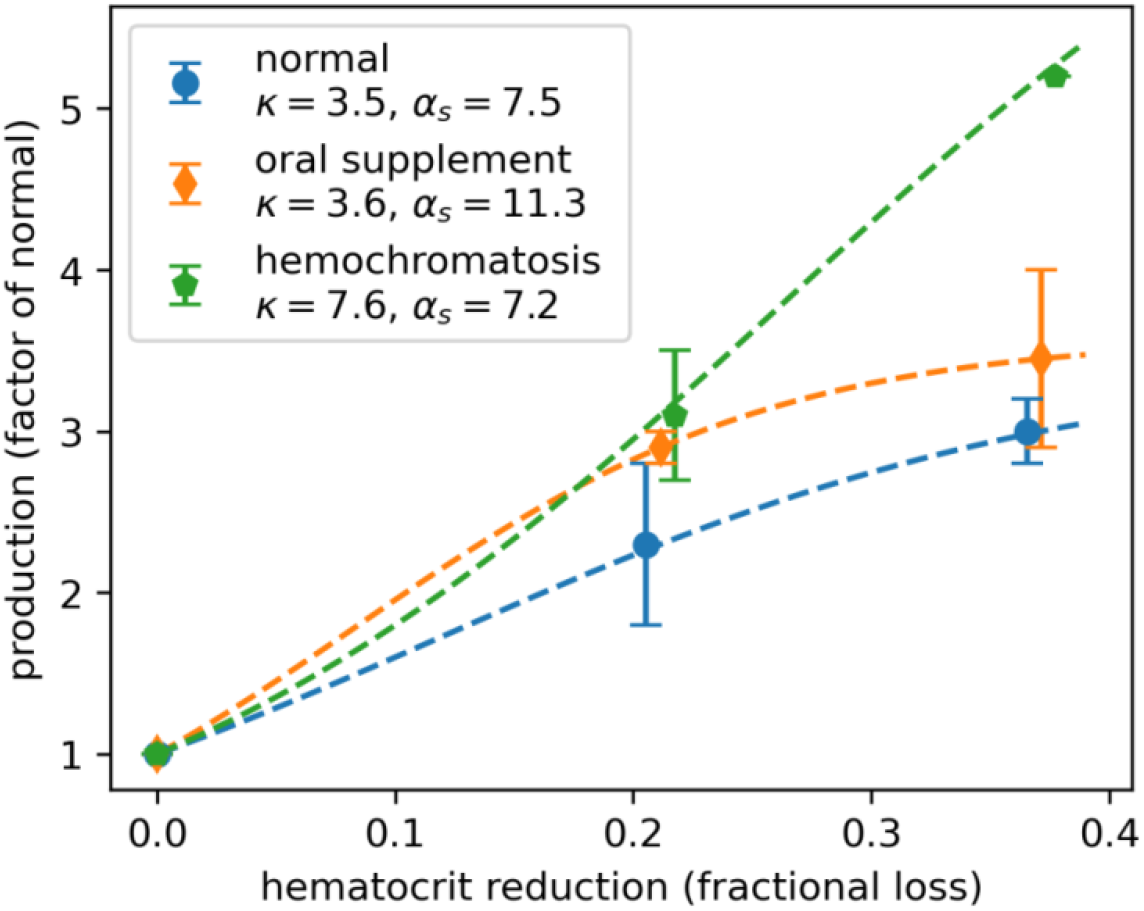
Parameter estimates based on Hillman et al. [20]. Three patient cohorts are defined by the authors based on the size of their available iron stores: a ‘normal’ control group, a group which was administered supplementary iron intakes, and a number of individuals suffering from hemochromatosis, a disorder characterized by an increased amount of total body iron stores. Each production factor shown (symbols) is the center of the range (error bars) measured within a patient cohort, as no individual measurements or averages are specified. Dashed lines result from a least-squares fit to the data employing the logistic coupling model, Eq. (3).

### Chronic perturbations lead to new equilibrium states

As a final exploration of the capabilities of the present model, we turn our attention to perturbations with a long-lasting character. These are of particular interest in medicine, as many genetic disorders such as inherited red cell membrane defects (hereditary spherocytosis, elliptocytosis, ovalocytosis), thalassemia syndromes and hemoglobinopathies (sickle cell disease, hemoglobin SC disease) all result in a chronic reduction of red cell survival times and anemia. Autoimmune hemolytic anemia due to autoantibodies against red blood cell antigens can also cause chronic destruction of red blood cells and anemia. We take as a model example the rare but well-studied paroxysmal nocturnal hemoglobinuria (PNH), a life-threatening disease characterized by an acquired mutation in the *PIGA* gene that renders red blood cells susceptible to complement attack resulting in severe hemolysis and other complications [24]. If the PNH afflicted clone is large enough, a significant portion of circulating erythrocytes will have a severely reduced lifespan. In our model we can take this into account by splitting the bloodstream compartment into a healthy (*H*) and a PNH afflicted (*PNH*) population, *N_B_* = *N_H_* + *N_PNH_*, where the death rate of the PNH group is significantly higher than that of the healthy cells (*μ_PNH_* > *μ_H_*). For a clone which comprises a fraction *p* of bone marrow cells, we obtain the dynamics

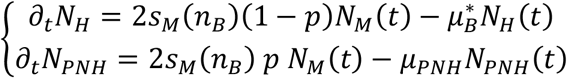

To determine which values of *p* might occur in humans, we note that PNH clones can comprise up to 100 percent of the blood cell population [25], while clones smaller than 10-20% could be considered subclinical. To obtain a realistic value for the rate at which these cells are hemolysed we use a study on the in-vivo survival rate of transfused erythrocytes from a PNH afflicted individual [26]. While no such death rate is derived in the paper itself, the authors describe a fast initial decay of the transfused population to 50% after only 5 days, followed by a slower decay down to 30% at the 10^th^ day. We can describe this behavior by means of two exponentially decaying populations to estimate the donor’s PNH fraction at *p* ≈ 0.8 and a death rate of *μ_PNH_* ≈ 0.2, which means that a PNH erythrocyte will be destroyed after on average 5 days in the bloodstream, 20 times faster than its normal counter-part.

Using the same parameter set derived in the previous section, we observe that, in the long term, new steady states emerge for all reacting compartments in any response regime (Fig 4). Using Fig 4a as a reference, we observe a marginal improvement in mitigating the loss in the self-renewal driven regime (*∂*_*n*_*v* < *∂*_*n*_*s*) (4b, 4e) whereas, in the differentiation driven regime (*∂*_*n*_*s* < *∂*_*n*_*v*) (4c, 4f) a reduced efficiency is observed. In contrast with the normal recovery that is realized under transient perturbations (Fig 2), the model also predicts a new stationary state for the bloodstream hemoglobin content, which in general remains below the normal homeostatic value. Furthermore, the model captures scenarios where the enduring reduced hemoglobin and red cell mass in circulation is accompanied by a persistent expansion of the upstream compartments (Figs 4b and 4e), as often seen in classic hemolytic PNH as well as other chronic hemolytic disorders [4]. As this expansion does not occur in the differentiation driven regime we conclude that the adaptive response in chronic hemolytic states must (at least) at times take place in a self-renewal driven regime.

**Figure 4.**
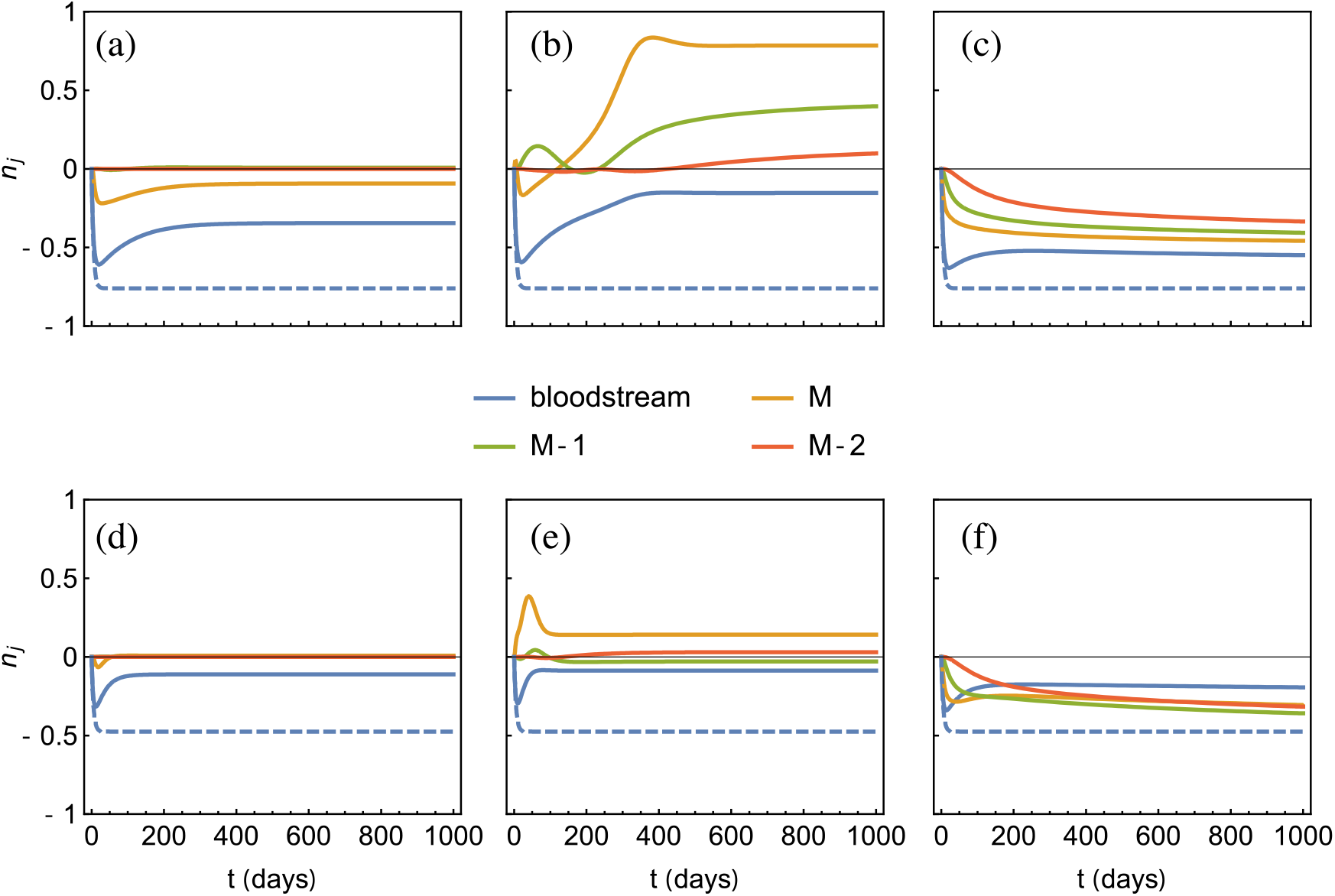
Dynamics following a chronic loss of cells in the bloodstream. Responses of an M=5 compartmental model employing logistic coupling with “normal” parameter values taken from Fig 3 (full lines) alongside the feedback-free response (dashed line). The rate of hemolysis of PNH afflicted erythrocytes is taken at μ_PNH_ = 0.2. Two different clone sizes are shown: p = 0.8 (panels (a)-(c)) and p = 0.5 (panels (d)-(f)). Balanced response to clone of size p is shown in panels (a) and (d), self-renewal driven response (*∂*_*n*_*v* < *∂*_*n*_*s*) to clone of size p is shown in panels (b) and (e), and differentiation driven response (*∂*_*n*_*s* < *∂*_*n*_*v*) to clone of size p is shown in panels (c) and (f).

## Discussion

The formalism described here provides a simple method for understanding the type of dynamics that populations of maturing hematopoietic cell precursors undergo in the bone marrow after being subject to different types of perturbations (from mild to severe), such as sudden or chronic blood loss. While the starting model of Dingli et al. [11] provides a useful framework for describing the hematopoietic system under homeostatic conditions, it does not account for the dynamics under perturbations such as those discussed here, as the time for a compartment to return to equilibrium is too long to fit clinically observed timescales [4]. The addition of sequential feedback to the model not only produces swifter recoveries, but also reproduces observed dynamic behaviors such as the response to a transient loss of erythrocytes, and the persistence of anemic states following chronic hemolysis with an associated chronic expansion of precursor cells in the bone marrow. The increased complexity, on the other hand, calls for a careful analysis of the properties of the feedback coupling introduced.

We identify three response types for any coupled pair of compartments, determined by the relative strengths of the differentiation and self-renewal coupling, *s*(*n*) and *v*(*n*) respectively. A perfectly balanced response prevents the perturbation from moving further upstream, thus providing the simplest reaction profile for hematopoiesis as a whole; it occurs whenever the equality *s*(*n*) – *v*(*n*) = *s*\* – *v*\* is fulfilled, and can intuitively be associated with a response where both differentiation and self-renewal increase (or decrease) in a balanced manner such that the compartment’s own cell number remains constant. This is however a very strict condition which is difficult to meet, even on average, in hematopoiesis, given its stochastic nature. Thus one expects that, in general, this detailed balance does not occur, and the dynamic behavior depends on which of the rates comes to dominate. When the differentiation rate dominates, the cell number in the compartment will change in the same direction as the perturbation – decreasing if the perturbation is a loss of cells, increasing if it is an excess – effectively introducing a positive feedback. When the self-renewal rate dominates, the compartment’s cell number varies in opposition with the perturbation – increasing with a loss, decreasing with an excess – which can lead to an overcompensation of the loss/excess followed by damped oscillations in the cell number. In some special cases where a resonant condition is met, nearly undamped oscillating cell counts in the blood are observed, associated with extreme cases found in certain hematologic disorders such as cyclic neutropenia [15,17].

It is important to take into consideration that in real hematopoiesis cell numbers in circulation fluctuate, even under homeostatic conditions [4]. Thus it is appropriate to introduce a range of values for the cell numbers within which hematopoiesis can be considered to be in (dynamic) equilibrium. In this sense small oscillations within this range predicted by our model can be presumed to be undetectable (and even if detectable, irrelevant) in a clinical setting. This in turn implies the rate parameters have some leeway to be out of sync without disturbing the bloodstream compartment in a detectable way, adding to the overall robustness of hematopoiesis. This feature, that leads to faster recovery times for small perturbations, may in turn result in long lasting perturbations for larger perturbations (Fig 2b, 2d). This highlights the importance of the stability of hematopoiesis with respect to the division rate parameters. Here, the coupling functions *s*(*n*) and *v*(*n*) posit a deterministic dependency of the division rates on downstream cell counts. In reality, these dependencies will be subject to noise from the underlying stochastic biological circuits and – as already pointed out – are unlikely to have perfectly balanced response solutions in the first place. Furthermore, since the response also depends on the availability of resources [20] which may vary or become depleted over time, the balance between s and *v* adaptation required for stability may itself change in time. However, an important observation is that this stability increases with increasing compartment number, or more specifically decreasing amplification between coupled compartments. The result furthermore adds an interesting angle to the currently favored view that normal hematopoiesis is mostly driven by ‘short-term’ stem cells which would be found further downstream then the small pool of long term HSCs [27,28], as such a larger pool of feedback coupled ‘drivers’ would increase stability.

An important quantifiable characteristic of the feedback driven system is the strength of the coupling between two compartments (determined by the values of the *α* and *k* parameters), as it governs the speed with which a return to equilibrium is attained. We find that while balanced responses (Fig 2a) allow for arbitrarily strong coupling, the physical limit of how fast a single cell can divide of course cannot be exceeded. Furthermore, the coupling strength may also depend on the availability of essential resources, as can be seen from a human erythropoiesis study where individuals with increased access to iron present amplified responses [20]. This observation raises the question of how long a particular response can be maintained, especially in the case of persistent losses.

Finally, it is worth remarking upon the differences between the compartmental dynamics under transient and chronic perturbations. In the former case, a short-lived perturbation such as bleeding can be swiftly remedied by increased cell divisions in the higher compartments, without propagating to earlier progenitor stages if the homeostatic balance between self-renewal and differentiation is maintained. In this sense the earliest compartments may not even be requested to respond to an acute loss of blood. On the other hand, chronic perturbations to the system – found in various hematopoietic disorders such as paroxysmal nocturnal hemoglobinuria and other hereditary or acquired hemolytic anemias – lead to the emergence of new equilibrium states that do not correspond to normal homeostasis. For example while the altered dynamics might mitigate a persistent loss of erythrocytes due to hemolysis by increasing the bone marrow output, the resulting steady-state number of erythrocytes in circulation may still be significantly lower than in the unperturbed system – a scenario which fits the observation of anemia occurring in severe cases of PNH as well as other hereditary or acquired hemolytic states. Experimental data from telomere length analysis in both PNH and sickle cell disease show that circulating mononuclear cells have shorter telomeres compared to age matched controls. Given that in our model self-renewal in any compartment is coupled to replication, this suggests that within hematopoiesis during chronic hemolysis, progenitor and downstream cells are undergoing more self-renewal — and thus more replication events than aged matched cells from healthy individuals, leading to shorter telomeres due to attrition with each replication [29,30]. In a former study [29] it was found that the shorter telomere length occurred in both PNH afflicted and unafflicted cells, suggesting that the cause indeed lies within hematopoiesis itself, suggesting that the feedback process intrinsic to hematopoiesis does not discriminate between the *PIGA* mutant and normal cells that co-exist in the bone marrow of patients with PNH.

## Acknowledgements

Nathaniel V. Mon Père gratefully acknowledges the funding of Télévie for supporting the research performed in this work.

## Supporting information

### S1 Original model and equilibrium values

In the model of Dingli et al. [11] the dynamics of a compartment *j* are given by

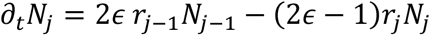

The first term on the right hand side of the equation is the flux of cells coming in from the nearest upstream compartment *j* – 1 (where the factor 2 comes from the fact that two daughter cells are created per division), while the second term is the sum of the fluxes of cells being removed due to differentiation (at rate *ϵr_j_N_j_*) and added due to self-renewal (at rate (1 – *ϵ*)*r_j_N_j_*). Given the number of HSCs *N*_0_, the number of (non-HSC) compartments *M,* and the daily bone marrow output *μ_M_*; the homeostatic values 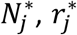, and *ϵ*\* can be found by simultaneously solving the equilibrium condition *η_ρ_* = 2*ϵ*(2*ϵ* – 1), the geometric growth equations *N_j_* = *N*_0_*η*^*j*^ and *r_j_* = *r*_0_*ρ^j^*, and the bone marrow output rate 2*ϵ r_M_N_M_* = *μ_M_* for *ϵ*, *η,* and *ρ*. For example, given a system with *M* = 28, *N*_0_ = 400, and *μ_M_* = 3.5 × 10^11^; the values *ϵ* = 0.82, *η* = 1.97, and *ρ* = 1.31 are obtained.

### S2 Competing Poisson processes

Consider the events *V* and *S* as respectively self-renewal and differentiation divisions, each with (independent) exponentially distributed waiting times with rates *v* and *s*. We have

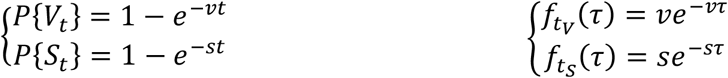

with *P*{*X_t_*} the probability of at least one event *X* occurring in time *t* and *f_t_x__*(*τ*) the density distribution of the waiting times. We now introduce the event 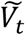 as the occurrence of at least one *V*, occurring before any *S* in *t*; and the complementary event 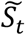 with the converse definition. Note that 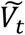 and 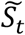 are mutually exclusive, and cover all possible outcomes except for those where no *S* or *V* occur in *t*. In our biological system these can be interpreted as two competing processes within the cell, where the first event to occur determines the divisional fate. We may write them equivalently as the following sets:

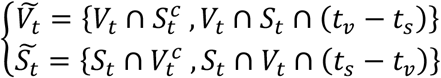

Since both events in each set are mutually exclusive we may write the probabilities of 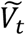 and 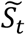 as the sum of the probabilities of their respective elements. The first term is

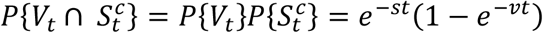

since *V_t_* and *S_t_* are independent (with the analogous argument for 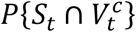). For the second term we obtain

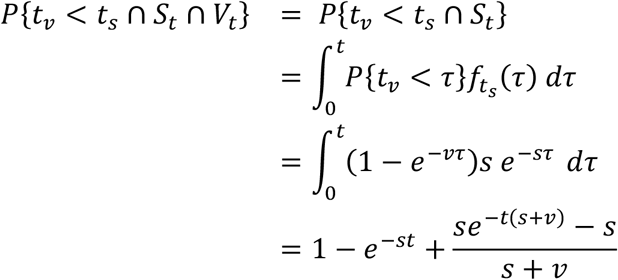

and summing the two gives

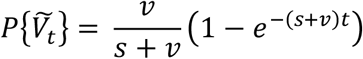

From this we identify (1 – *e*^−(*s*+*v*)*t*^) = *P*{*V_t_* ∪ *S_t_*}, which is the probability of any division (self-renewal or differentiation) occurring in *t*. Thus 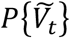 and 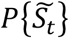 can readily be interpreted as the probabilities of a division occurring in t, multiplied by a probability which determines whether that division is a self-renewal or a differentiation. The above expression can furthermore be expanded for an infinitesimal timestep *dt* to obtain a rate [31]:

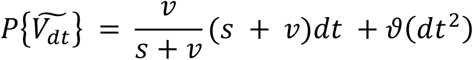

which makes it clear that differentiation and self-renewal occur at rates *s* and *v* respectively, and allows us to identify *ϵ* = *s*/(*s* + *v*) and *r* = *s* + *v*.

## References

1. Doulatov S, Notta F, Laurenti E, Dick JE. Hematopoiesis: A Human Perspective. Cell Stem Cell. 2012;10: 120–136. doi:10.1016/j.stem.2012.01.006

2. Laurenti E, Göttgens B. From haematopoietic stem cells to complex differentiation landscapes. Nature. 2018;553: 418–426. doi:10.1038/nature25022

3. Notta F, Zandi S, Takayama N, Dobson S, Gan OI, Wilson G, et al. Distinct routes of lineage development reshape the human blood hierarchy across ontogeny. Science. 2016;351: aab2116. doi:10.1126/science.aab2116

4. Kaushansky K, editor. Williams hematology. Ninth edition. New York: McGraw-Hill; 2016.

5. Eaves CJ. Hematopoietic stem cells: concepts, definitions, and the new reality. Blood. 2015;125: 2605–2613. doi:10.1182/blood-2014-12-570200

6. Höfer T, Rodewald H-R. Differentiation-based model of hematopoietic stem cell functions and lineage pathways. Blood. 2018;132: 1106–1113. doi:10.1182/blood-2018-03-791517

7. Robb L. Cytokine receptors and hematopoietic differentiation. Oncogene. 2007;26: 6715–6723. doi:10.1038/sj.onc.1210756

8. Bélair J, Mackey MC, Mahaffy JM. Age-structured and two-delay models for erythropoiesis. Mathematical Biosciences. 1995;128: 317–346. doi:10.1016/0025-5564(94)00078-E

9. Bernard S, Bélair J, Mackey MC. Oscillations in cyclical neutropenia: new evidence based on mathematical modeling. Journal of Theoretical Biology. 2003;223: 283–298. doi:10.1016/S0022-5193(03)00090-0

10. Adimy M, Crauste F, Ruan S. A Mathematical Study of the Hematopoiesis Process with Applications to Chronic Myelogenous Leukemia. SIAM J Appl Math. 2005;65: 1328–1352. doi:10.1137/040604698

11. Dingli D, Traulsen A, Pacheco JM. Compartmental Architecture and Dynamics of Hematopoiesis. PLOS ONE. 2007;2: e345. doi:10.1371/journal.pone.0000345

12. Marciniak-Czochra A, Stiehl T, Ho AD, Jäger W, Wagner W. Modeling of Asymmetric Cell Division in Hematopoietic Stem Cells—Regulation of Self-Renewal Is Essential for Efficient Repopulation. Stem Cells and Development. 2008;18: 377–386. doi:10.1089/scd.2008.0143

13. Lo W-C, Chou C-S, Gokoffski KK, Wan FY-M, Lander AD, Calof AL, et al. FEEDBACK REGULATION IN MULTISTAGE CELL LINEAGES. Math Biosci Eng. 2009;6: 59–82. doi:10.3934/mbe.2009.6.59

14. Doumic M, Marciniak-Czochra A, Perthame B, Zubelli J. A Structured Population Model of Cell Differentiation. SIAM J Appl Math. 2011;71: 1918–1940. doi:10.1137/100816584

15. Pacheco JM, Traulsen A, Antal T, Dingli D. Cyclic neutropenia in mammals. American Journal of Hematology. 2008;83: 920–921. doi:10.1002/ajh.21295

16. Dingli D, Traulsen A, Pacheco JM. Dynamics of haemopoiesis across mammals. Proceedings of the Royal Society of London B: Biological Sciences. 2008;275: 2389–2392. doi:10.1098/rspb.2008.0506

17. Dingli D, Antal T, Traulsen A, Pacheco JM. Progenitor cell self-renewal and cyclic neutropenia. Cell Proliferation. 2009;42: 330–338. doi:10.1111/j.1365-2184.2009.00598.x

18. Lenaerts T, Pacheco JM, Traulsen A, Dingli D. Tyrosine kinase inhibitor therapy can cure chronic myeloid leukemia without hitting leukemic stem cells. Haematologica. 2010;95: 900–907. doi:10.3324/haematol.2009.015271

19. Mon Père N, Lenaerts T, Pacheco JM, Dingli D. Evolutionary dynamics of paroxysmal nocturnal hemoglobinuria. PLOS Computational Biology. 2018;14: e1006133. doi:10.1371/journal.pcbi.1006133

20. Hillman RS, Henderson PA. Control of marrow production by the level of iron supply. J Clin Invest. 1969;48: 454–460. doi:10.1172/JCI106002

21. Otto SP, Day T. A biologist’s guide to mathematical modeling in ecology and evolution. Princeton: Princeton University Press; 2007.

22. Strogatz SH. Nonlinear dynamics and chaos: with applications to physics, biology, chemistry, and engineering. 2. print. Cambridge, Mass: Perseus Books; 2001.

23. Werner B, Dingli D, Lenaerts T, Pacheco JM, Traulsen A. Dynamics of Mutant Cells in Hierarchical Organized Tissues. PLOS Computational Biology. 2011;7: e1002290. doi:10.1371/journal.pcbi.1002290

24. Brodsky RA. Paroxysmal nocturnal hemoglobinuria. Blood. 2014;124: 2804–2811. doi:10.1182/blood-2014-02-522128

25. Socié G, Schrezenmeier H, Muus P, Lisukov I, Röth A, Kulasekararaj A, et al. Changing prognosis in paroxysmal nocturnal haemoglobinuria disease subcategories: an analysis of the International PNH Registry. Internal Medicine Journal. 2016;46: 1044–1053. doi:10.1111/imj.13160

26. Dacie JV, Mollison PL. SURVIVAL OF TRANSFUSED ERYTHROCYTES FROM A DONOR WITH NOCTURNAL HÆMOGLOBINURIA. The Lancet. 1949;253: 390–392. doi:10.1016/S0140-6736(49)90704-7

27. Sun J, Ramos A, Chapman B, Johnnidis JB, Le L, Ho Y-J, et al. Clonal dynamics of native haematopoiesis. Nature. 2014;514: 322–327. doi:10.1038/nature13824

28. Busch K, Klapproth K, Barile M, Flossdorf M, Holland-Letz T, Schlenner SM, et al. Fundamental properties of unperturbed haematopoiesis from stem cells *in vivo*. Nature. 2015;518: 542–546. doi:10.1038/nature14242

29. Karadimitris A, Araten DJ, Luzzatto L, Notaro R. Severe telomere shortening in patients with paroxysmal nocturnal hemoglobinuria affects both GPI− and GPI+ hematopoiesis. Blood. 2003;102: 514–516. doi:10.1182/blood-2003-01-0128

30. Mekontso Dessap A, Cecchini J, Chaar V, Marcos E, Habibi A, Bartolucci P, et al. Telomere attrition in sickle cell anemia. American Journal of Hematology. 2017;92: E112–E114. doi:10.1002/ajh.24721

31. Feller W. An introduction to probability theory and its applications. Vol. 1. 3. ed., rev. print., [Nachdr.]. S.l.: Wiley; 2009.

